# The consequences of mismatched buffers in spectral cell sorting

**DOI:** 10.1101/2024.08.19.608560

**Authors:** Rita A. S. Dapaah, Laura Ferrer-Font, Xiaoshan Shi, Christopher Hall, Sam Thompson, Larissa Catharina Costa, Peter L. Mage, Aaron J. Tyznik, Kelly Lundsten, Rachael V Walker

## Abstract

Although spectral flow cytometry has become a ubiquitous tool for cell analysis, the use of spectral cytometry on cell sorters requires additional considerations arising from the unique requirements of sorting workflows. Here, we show that care should be taken when ascertaining the purity of a sort on a spectral cell sorter, as the mismatch of buffers used for initial sample suspension and the buffers used for sort collection can affect the unmixing of the data, potentially giving rise to erroneous purity check results.

## INTRODUCTION: Purity check observations

Cell sorting was first established in 1965[1] by Mack Fulwyler, where the determination of a successful sort was assessed by measuring the enrichment of cell concentrations after sorting when compared to the pre-sorted population. This method of sort validation continued in the Hulett *et al* Science paper[2] where sorted Chinese hamster ovary (CHO) cells were checked using volumetric and fluorescence intensity measurements. From the inception of cell sorting, an assessment of the sorted cells has been used to judge the success of the sort (a purity check).

Advances in flow cytometry hardware have resulted in the release of multiple spectral flow cytometer cell sorters, including the Thermo Fisher Invitrogen Bigfoot (Bigfoot), the BD FACSDiscover™ S8 Cell Sorter (BD FACSDiscover™ S8), and the Cytek Biosciences Aurora CS system (CS). Spectral flow cytometry uses the signal from all available fluorescence detectors to identify unique fluorescent signatures compared to conventional cytometry that relies upon the peak emission detector to define the presence of a fluorochrome and corresponding antibody on the cells of interest. [3] [4] [5] This requires different signal deconvolution methodology, colloquially termed spectral unmixing where peak emission-based analysis uses an approached referred to as “compensation.”

In this technical note, we report post-sort purity checks from BD FACSDiscover™ S8, Aurora CS, and Bigfoot spectral sorters where sorted populations fell outside of the software defined sort gates upon reanalysis (Figure 1). Although these results initially led us to call into question the success of each sort, we realized that the observed purity check could also be explained by a change in the spectrally unmixed signal of the sorted sample. Here we explore these observations and investigate possible causes of a change in spectral signal between pre- and post-sorted samples, ultimately concluding that spectral shifts arising from pre- and post-sort buffer conditions are the likely cause.

**Figure 1:**
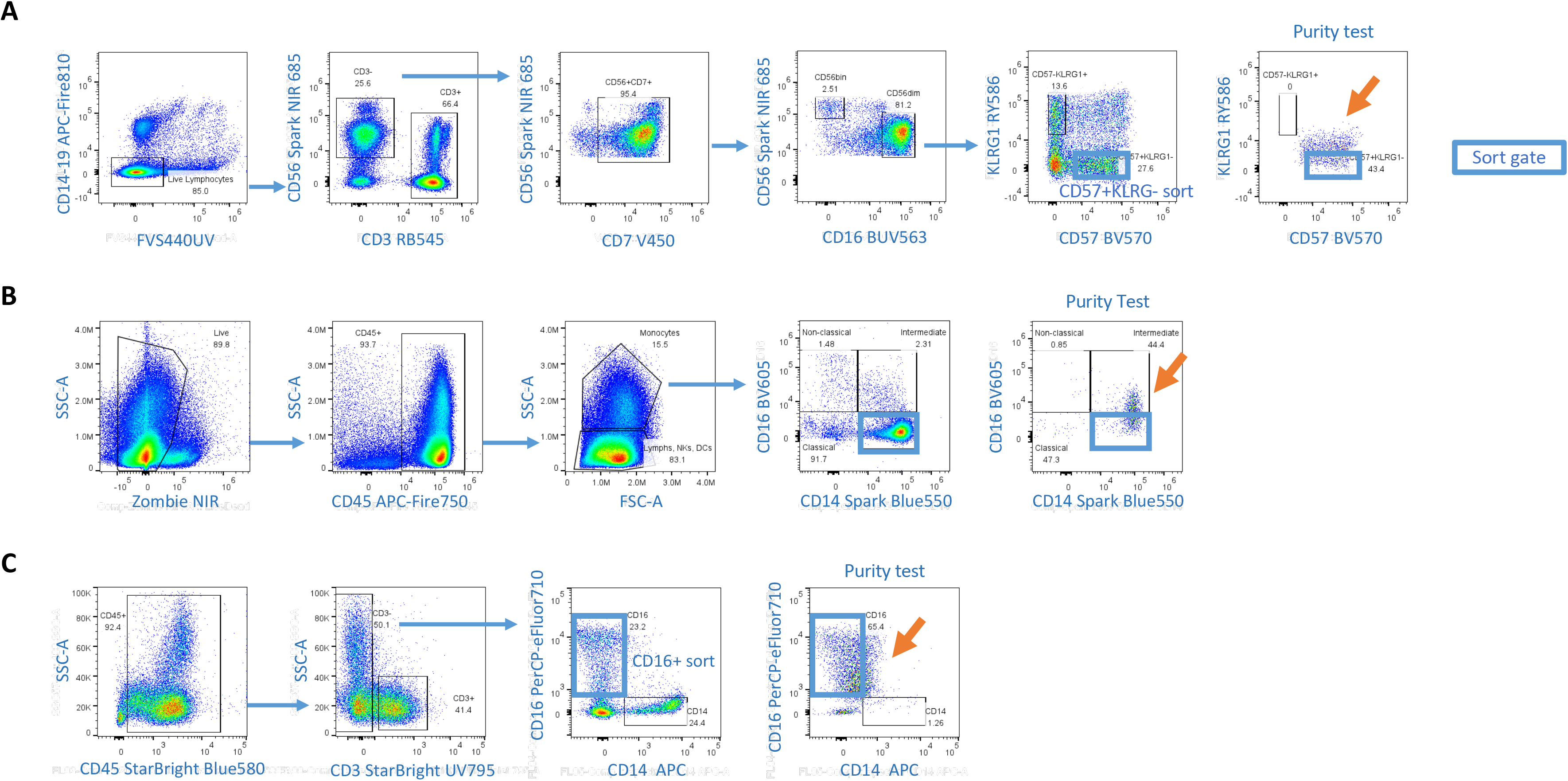
Population shifts are visualised post-sort on 3 instruments and 3 different panels. Solid blue outline indicates the population sorted and orange arrow indicates the post-sort purity check population shift. (**A**) 9 fluorescent marker panel on the BD FACSDiscover™ S8, CD57+ cells sorted and re-acquired. (**B**) 26 fluorescent marker panel on Cytek Biosciences Aurora CS, singlets cells sorted and re-acquired. (**C**) 6 fluorescent marker panel on the Thermo Fisher Invitrogen Bigfoot Cell Sorter, CD16+ cells sorted and re-acquired.

### Hypothesis 1: Evaluation of spectral shifts post sort

We initially theorized that the spectral shift observed in sorted cells was caused by the sorting process itself, which not only exposes the cell-bound fluorochromes to potential photobleaching from laser illumination, but also exposes the cells to varying pressure conditions throughout the sample fluidics, nozzle, and sort block. To evaluate spectral shifts that may arise from these factors, the single-stain controls and fully stained sample of a six-fluorophore panel were sorted on the Bigfoot sorter three sequential times with the evaluation of the resulting spectra for each sample (pre-sort, 1^st^ sort, 2^nd^ sort, and 3^rd^ sort). The pre-sort sample and post-sort collection buffers were identical. Supplementary Figure 1 shows this experiment, with the gating scheme and each purity check overlaid (1a). Figure 1b shows the spectra of the fully stained sample as pre-sort and each round of sorting plus the similarity matrix. Figure 1c shows each individual fluorophore and its similarity matrix. No spectral changes were seen in the similarity matrix suggesting that spectral changes from repetitive sorts did not have a significant role in the observed poor purities previously observed where the samples passed through the sorter once.

### Hypothesis 2: Effects of buffer changes

We next hypothesized that the pre- and post-sort buffer conditions may be a cause of spectral differences upon sorting. The buffer that cells are deposited into will often vary depending on the cell type and downstream processing required. It was noted that during the cell sorts and ensuing purity checks, a common feature was that the initial resuspension buffer for that specific sort application was different than the collection buffer that the cells were resuspended in during the purity check. To evaluate the potential impact of buffers on post-sort purity checks, we performed a series of experiments to evaluate the impacts of different buffers and the visualization of the cell populations on user-defined gated populations.

On the spectral instruments evaluated, it was observed that the buffer into which the cells are sorted impacts the perceived purity measured. A 9-fluorophore panel was acquired on the BD FACSDiscover™ S8 (Figure 2A, B, C, and F), on the Aurora (Figure 2 D, E and G), and the Aurora CS (Figure 2H) using PBS+BSA buffer for the single colour unmixing controls and as the reference standard. The CD57+KLRG1-population shift was observed when running the post-sort purity check into BD FACSFlow™ (Figure 2A). Initial observations would suggest that there is an unmixing error derived from BUV563 (CD16) in (Figure 2B) and the signature overlay of a single stain CD16 BUV563 shows a small difference in the spectral signature that may be the cause of the unmixing issue (Figure 2C). To further explore the impact of the buffers on resultant purity checks, we distributed the same fully stained sample into different buffer conditions and confirmed apparent spectral shifts by utilizing different instruments for independent analysis (Figure 2F-H). Figure 2E shows the signature overlay of a single stain CD16 BUV563 on the Aurora analyser, the spectral change is not as pronounced as on the S8 (figure 2C) probably due to different filters sets between the instruments and different buffers evaluated. To confirm the observed shifts in populations came from different buffer conditions, first the fully stained sample was resuspended in PBS+BSA Buffer and acquired on the BD FACSDiscover™ S8 (Figure 2F). The same sample was then spun down and resuspended in BD FACSFlow™ followed by acquisition on the same instrument. Finally, the sample was spun down again and resuspended back into PBS+BSA buffer and re-acquired. Similar to post-sort analysis in Figure 2A, using the unmixing matrix compiled from the PBS+BSA single colour controls, there is an observed shift of the all KLRG1-cells whether they are CD57 BV570 positive or negative in this plot along the RY586 axis on the bivariate plot in the BD FACSFlow™ condition. However, the subsequent resuspension back into the original PBS+BSA buffer reversed this effect, placing the original populations in the gate as initially observed under the same condition (Figure 2F). To confirm that these results are not specific to one instrument or a specific combination of buffers, a similar set of experiments were performed using both an Aurora Analyser (Figure 2G) and Aurora CS (Figure 2H). For these experiments, different buffers were evaluated, cells were initially resuspended with Biolegend staining buffer (contains BSA) and single stained controls in the staining buffer were used for unmixing, followed by acquisition of the full panel. The same sample was then spun down and resuspended in BD FACSFlow™ Buffer and acquired with the initial unmixing matrix (Figure 2G and 2H). In this example shifts in populations were also observed using different buffers for resuspension compared to original buffer, highlighting the impact of the resuspension buffer when evaluating sort purity.

**Figure 2.**
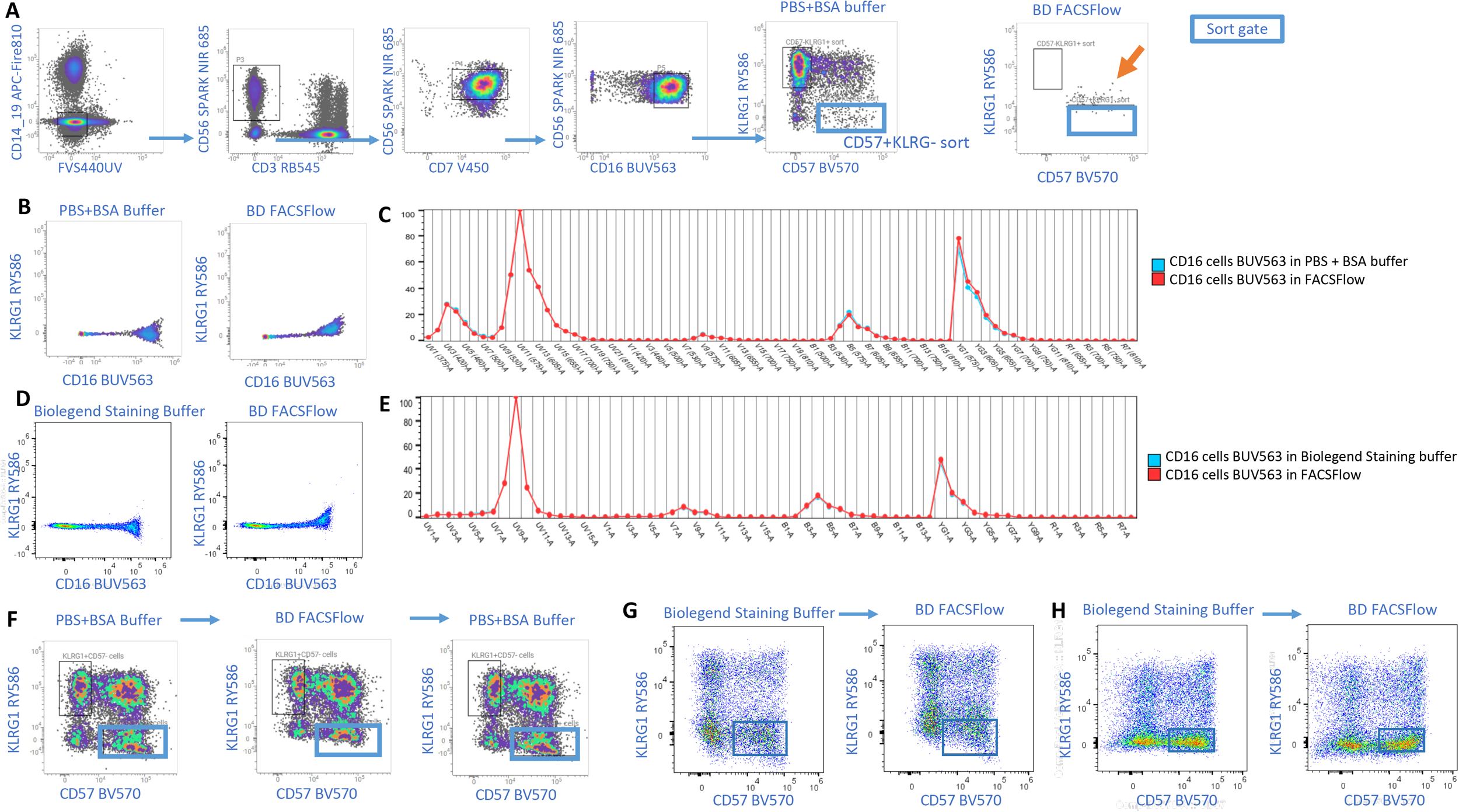
9 fluorescent marker panel on the BD FACSDiscover™ S8, Cytek Biosciences Aurora analyser and Aurora CS. (**A**) Gating strategy of a full stained sample in PBS+BSA buffer unmixed with single stain cells controls in PBS+BSA buffer. The sorted population gate is marked in a blue square with an ultimate marker phenotype of live/CD14-/CD56+/CD7+/CD16+/KLRG-. Cells were sorted into BD FACSFlow™ (no added protein) and re-acquired on the BD FACSDiscover™ S8 where a population shift can be observed in the CD57+/KLRG1-sorting gate (blue square, far right plot). (**B**) CD16 BUV563 Single stain cells resuspended in PBS+BSA buffer and BD FACSFlow™ on the BD FACSDiscover™ S8. (**C**) Spectral signature overlays of CD16 BUV563 Single Stain Cells resuspended in PBS+BSA buffer (blue line) and BD FACSFlow™ (red line) on the BD FACS Discover™ S8. (**D**) CD16 BUV563 Single Stain Cells resuspended in Biolegend Staining Buffer (contains BSA) and BD FACSFlow™ on Cytek Aurora Analyser. **(E)** Spectral signature overlays of CD16 BUV563 Single Stain Cells resuspended in Staining Buffer (blue line) and BD FACSFlow™ (red line) on Cytek Aurora Analyser. (**F**) Full stained sample in PBS+BSA, washed and resuspended in BD FACSFlow™ and washed and resuspended into PBS+BSA and acquired on the BD FACSDiscover S8 (these samples were not sorted only acquired; blue box indicates population of interest) (**G**) Full stained sample resuspended in Biolegend staining buffer, washed, and resuspended in BD FACSFlow™ and acquired on the Aurora Analyser (blue box indicates population of interest). (**H**) Full stained sample resuspended in Biolegend staining buffer, washed, and resuspended in BD FACSFlow™ and acquired on the Aurora CS (these samples were not sorted only acquired, blue box indicates population of interest). All cells used in this figure were unfixed apart from figure 2H where the cells were fixed with Biolegend Fixation buffer due to the biosafety regulations.

To confirm that this buffer-dependent spectral shift is not specific to a single particular reagent, additional panels were evaluated on the Aurora CS and Bigfoot sorters which confirmed that a phenomenon similar to that observed in Figure 1 and 2 occurs for additional reagents and panels (Supplementary Figure 2).

To further define the impact of buffers on a fully stained panel, we performed a series of experiments to confirm that this effect is not specific to cells being used as single stained controls. The use of single stained compensation beads for unmixing instead of cells was evaluated, using the same panel as described in Figure 2. Supplementary Figure 3A, shows the single stained compensation beads stained in PBS+BSA and unmixed on the BD FACSDiscover™ S8. When the fully stained sample was subsequently re-suspended and re-acquired in BD FACS Pre-Sort buffer a shift in the CD57+ and CD57-cells was observed. To confirm that the BD FACS™ Pre-sort buffer was not the cause of the problems with the panel, single stained beads were resuspended into the BD FACS™ Pre-Sort buffer and acquired on the FACSDiscover™ S8 and unmixing carried out. The fully stained sample resuspended also in the BD FACS™ Pre-Sort buffer was then acquired and no shift in populations was seen, showing that if the buffers are matched, no shift in populations are observed (Supplementary Figure 3B). Supplementary Figure 3C shows an overlay of single stained CD16 BUV563 beads resuspended in PBS+BSA and BD FACS™ Pre-Sort buffer. The difference in spectral signatures can be found and are very similar to those seen to the single cell overlay in Figure 2.

As part of the study, autofluorescence (AF) changes in samples during sorting and the different buffers were investigated. As observed in Supplementary Figure 4, the addition of different buffers can have a clear impact on the spectral signature of the unstained samples. Importantly, if samples are matched with an unstained in the same resuspension media, unmixing issues can be reduced. In the case of Figure 2 in which a population shift was observed, it was important to ensure that the AF was not the cause of the population shift. Two unmixing matrices were created, one with AF extraction and one without AF extraction. Data was compared (Supplementary Figure 5) and the AF extraction did not influence the population shift.

## CONCLUSIONS

The study has shown that spectral cytometry and especially spectral cell sorting has revealed the complex chemistry that can occur when fluorophores are exposed to different buffers. Although we did not explore the exact underlying mechanism behind the observed results, it is possible that these spectral changes could be caused by changes in the fluorophores’ chemical microenvironment due to ions and pH in buffers, various additives that may have their own spectral profiles, as well as sources of changes that have yet to be defined. These changes may not be predictable as seen in the results when the samples were run on different spectral sorters. It should be noted that these changes also occur in fluorochromes often used in conventional cytometry and cell sorting. While evaluating the emission profiles, the changes seen in spectral signatures are often in areas of the spectra that are located away from the peak channel but may still affect the unmixing of the samples. These subtle changes in each fluorochrome may be compounded when multiple fluorochromes are being used together resulting in poor purity checks. As the number of parameters in spectral cytometry is constantly increasing (recently published OMIP-102 shows a 50 fluorescent parameter panel[6]), these changes are likely to have increasing problems with larger and more complex panels.

To summarize the observation, analysing cells in one suspension buffer and sorting into another can lead to changes in the fluorochrome emission and/or excitation profiles. These changes may result in a mismatch in the spectral signature of sorted cells and single-stain controls that cause unmixing errors. Upon post-sort purity checks, there may be a population shift suggesting a potentially failed cell sort. This study has shown that spectral changes can occur under different buffer conditions resulting in inaccurate conclusions about a sort. We believe these results are of note for anyone in the spectral cytometry community aiming to achieve reliable and reproducible sorting results.

## METHODS

### Evaluation of spectral shifts post sort

Frozen mouse splenocytes (mix of *Fcer2a*^cre/+^; *Cd69*^fl/fl^ and *Fcer2a*^+/+^;*Cd69*^fl/fl^ cells) were thawed in a 37°C water bath for 2 mins and resuspended in 10 ml of warm Staining Buffer (SB; Biolegend #420201). Cells were washed twice at 300 g for 10 mins. Cells were resuspended at 10 × 10^7^/mL in SB. A master mix (5 µL of each fluorochrome-conjugated antibody (list of antibodies in supplementary table 1) + 10 µL of Brilliant Staining Buffer (BD Horizon™ #659611) was added to 25-50 × 10^6^ cells in 250 µL of SB. For single colour controls and 2-5 × 10^6^ cells were added to 5 µL of fluorochrome-conjugated antibody. Cells were incubated in the dark at 4°C for 45 minutes. Cells were washed twice with 2 mL of SB (300 g for 5 mins). Single colour controls were resuspended in 200 µL of SB and full stain sample were resuspended at 10 × 10^6^/mL in SB.

All cells were sorted using a singlet gate (FSC-A vs SSC-A and FSC-H vs SSC-H) into staining buffer. The sorted sample in their entirety were then sorted through the Bigfoot again twice more. A final purity check of an aliquot of cells was carried out. Unmixing was carried out using Sasquatch Software (SQS) v.1.19.41805.4 and data analysis done using FlowJo v.10 software.

### Effects of buffer changes

#### Cell preparation and staining

Frozen PBMCs and fresh PBMCs were used. Frozen human peripheral blood mononuclear cells (PBMCs) were thawed in a 37°C water bath for 2 mins and resuspended in 10 ml of warm staining buffer (SB; Biolegend #420201) or media buffer (Thermo Fisher Scientific DMEM #12634010 or RPMI #61870010) with 20% Foetal Calf Serum (FCS, Gibco, A380001). Cells were washed twice at 300 g for 10 mins. Cells were resuspended in SB or 1X Phosphate-buffered saline (PBS).

#### 9 fluorescent marker panel

Figures 1A and 2, supplementary 3 and supplementary 5: Thawed frozen PBMCs (work carried at Babraham Institute), fresh PBMCs (work carried out at BD Life Sciences-Bioscience) were resuspended at 1 × 10^6^ in 100 µL of PBS for single colour controls and 5 × 10^6^ in 100 µL of PBS for full stain sample. For the live-dead control 50µL of cells were heat shocked (65 ^⍰^C for 5 min), resuspended with 50µL of live cells. Sample and live-dead control were treated (/stained) with 1:1000 fixable viability stain (FVS) 440UV at 37^⍰^C for 10 min. Single colour controls and full stain sample were added to 10 µL Brilliant Staining Buffer Plus (BD Horizon™ # 568264) and fluorochrome-conjugated antibodies at indicated concentrations (Supplementary Table 1) and incubated at Room temperature (RT) for 30 mins. Cells were washed twice with 2 mL of PBS + BSA (400 g for 5 min) and run fixed (Biolegend fixation buffer) or unfixed prior to sorting and analysis. Cells were resuspended in PBS + BSA or BD FACSFlow™ (BD, 342003). For supplementary figure 3, compensation beads (BD Biosciences) were stained in the same way as cells for single colour controls and used for spectral unmixing.

#### 26 fluorescent marker panel – Figure 1B

Thawed frozen PMBCs (50 × 106 cells) and an aliquot of heat-treated cells for viability single control (2 × 106) were respectively resuspended in 500 and 100 µL of Zombie NIR™ solution (1:2000) and incubated at RT for 15-30mins in the dark. Cells were washed twice with 2 mL SB (300 g for 5 mins) and resuspended in 225 µL of SB. Cells were added to 25 µL Human TruStain FcX™ (Biolegend #422301) and incubated at RT for 10 mins. Single colour controls (2 × 10^6^ cells in 100 µL SB) and full stain sample were stained with fluorochrome-conjugated antibodies at indicated concentrations (Supplementary Table 1) and incubated at 4°C for 30 mins. Cells were washed with 2 mL SB (350 g for 5 min) and subsequently with 2 mL of 1X Tandem Stabilizer (Biolegend #421802). Cells were resuspended in 200 µL of 1X Tandem Stabilizer.

#### Figures 1c and Supplementary 1 and supplementary figure 2

Thawed PBMCs were resuspended at 10 × 10^7^/mL in SB (Biolegend #420201). A master mix (5 µL of each fluorochrome-conjugated antibody + 10 µL of Brilliant Staining Buffer (Figure 1 a & b) (BD Horizon™ #659611) or Super Bright Complete Staining Buffer, (figure 1c) (eBioscience #SB-4401-42)) was added to 25-50 × 10^6^ cells in 250 µL of SB. For single colour controls and 2-5 × 10^6^ cells were added to 5 µL of fluorochrome-conjugated antibody (see supplementary table 1 for fluorochrome details). Cells were incubated in the dark at 4°C for 45 minutes. Cells were washed twice with 2 mL of SB (300 g for 5 mins). Human cells were fixed Biolegend fixation buffer (4% Paraformaldehyde) prior to analysis and sorting. Single colour controls were resuspended in 200µL of SB and full stain sample were resuspended at 10 × 10^6^/mL in SB.

#### Unstained cells - Supplementary 4

*a & b)* Veri-Cells™ (Biolegend #425001) were reconstituted following manufacturer guidelines. Lyophilized cells were warmed at RT for 5 mins, reconstituted with Veri-Cells™ Buffer A Plus and left at RT for 15 mins prior to staining. Cells were centrifuged at 300g for 5 mins and resuspended in 150µL (96 U-well plates) or 300µL (5ml tubes) of different buffer solution. *c)* Frozen human PBMCs were thawed and fixed as above and resuspended in 150µL (96 well plates) or 300µL (5ml tubes) of the different buffer solutions. d) Frozen mouse splenocytes were thawed and fixed as above and resuspended in 150µL (96 well plates) or 300µL (5ml tubes) of the different buffer solutions.

Different buffer solutions used in supplementary figure 4: Phosphate Buffered solution (PBS) (in house (Sodium Chloride, Di-Sodium Hydrogen Phosphate Dihydrate)); PBS with 10% Foetal Calf Serum (FCS) (Gibco, A380001); BD FACSFlow™ (BD, 342003)); Dulbecco’s Modified Eagle’s Medium (DMEM) without Phenol Red (Thermo Fisher, 11880028) and 10% FCS; DMEM with Phenol Red (Thermo Fisher Scientific, 12634010) with 10% FCS; RPMI with Phenol Red (61870010) and 10% FCS; Hank’s buffered solution (H6648) with 10% FCS.

##### Data unmixing and data analysis

Data were compensated (BD FACSDiva™ v.8.0.1 Software) and unmixed (BD FACSChorus™ v5.1 Software, Sony™ ID700 v.2.0.1, SpectroFlo™ v.3.2.1, SpectroFlo™ CS v.1.0.7 and Sasquatch Software (SQS) v.1.19.41805.4) using the respective acquisition software for each instrument, whilst spectral signatures normalization and comparisons were performed in FlowJo™ v.10 Software or R studio v.4.3.1. Data files in Flow Repository show a reduced number of 20,000 events for each file acquired.

### Instrument details

All samples were run on standard Cytek Aurora 5 laser analyser, Cytek Aurora CS; BD FACSDiscover™ S8 and Thermo Fisher Invitrogen Bigfoot. Laser and filter configurations can be found in Supplementary Table 2.

## ACKNOWLEDGEMENTS

Babraham Institute would like to acknowledge the Babraham Institute Flow Cytometry Facility for their assistance with this project, Dr Oliver Burton for reviewing this manuscript, Dr Jonathan Clark for discussions, and Stephane Guillaume for providing the mouse splenocytes. The Babraham Institute Flow Facility is funded by the BBSRC Core Capability Grant. Human work carried out at Babraham Institute is covered by ethical approval (23/NW/0287). All experiments using human whole blood from healthy donors were carried out in compliance with the criteria of the Environment Health and Safety office in BD Biosciences.

Christopher Hall and Laura Ferrer-Font are current ISAC Emerging Leaders.

Rachael Walker is a past ISAC Emerging Leader

Peter Mage is a current ISAC International Innovator

Aaron Tyznik is a past ISAC MaryLou Ingram Scholar

## Figures and tables

**Supplementary Figure 1: 6 fluorescent marker panel (mouse) sorted on the Thermo Fisher Invitrogen Bigfoot, the same sample was sorted three times using the singlets gate**. (A) Gating strategy of full stain sample pre-sort overlaid with the 3 subsequent sort. (B) Full panel normalised spectrum, pre-sort and 3 subsequent sort overlaid and the similarity matrix. (C) Single colour controls normalised spectrum, pre-sort and 3 subsequent sort overlaid and the similarity matrix

**Supplementary figure 2: 6 fluorescent marker panel (human) gating strategy and purity test on Cytek Biosciences Aurora CS and Thermo Fisher Invitrogen Bigfoot**. Gating strategy of a full stained sample in Biolegend Staining buffer unmixed with single stain cells controls in Biolegend Staining buffer. CD16+ population were sorted on the (**A**) Cytek Biosciences Aurora CS and (**D**) Thermo Fisher Invitrogen Bigfoot. CD16+ cells were sorted into staining buffer (control, B and E) and DMEM with 10% FCS (**C** and **F**) and re-acquired on the sorters. A population shift can be observed for the cells sorted into and DMEM with 10% FCS. An aliquot of CD16+ cells sorted into DMEM with 10% FCS were then washed and resuspended into Biolegend staining buffer and reacquired on the sorters (**C** and **F**).

**Supplementary figure 3. 9-fluorescent marker panel run on the BD FACSDiscover** ^**TM**^ **S8 using single stain beads. (A)** Full stained (FS) sample in PBS+BSA buffer unmixed with single stain (SS) beads controls in PBS+BSA buffer, the FS was washed and resuspended in BD FACS™ Pre-sort buffer. **(B)** SS controls were resuspended in Pre-sort buffer and used to unmix FS sample resuspended in pre-sort buffer. **(C)** Spectral signature overlays of CD16 BUV563 Single Stain Beads resuspended in PBS+BSA buffer (blue) and Pre-sort buffer (red).

**Supplementary figure 4: Difference in autofluorescence observed across a variety of cell types in different buffer solutions**. Normalised spectral signatures of unstained cells in different buffer overlaid. (**A-B**) Veri-Cells PBMCs (lymphocytes and monocytes) resuspended in six buffer solutions (PBS, BD FACSFlow™, DMEM without phenol and 10% FCS, DMEM with phenol and 10% FCS, RPMI with phenol and 10% FCS, HANKS and 10% FCS) and (**B-C**) fixed human PBMCs and mouse splenocytes resuspended in seven buffer solutions (BD FACSFlow™, DMEM with phenol, DMEM with phenol and 10% FCS, DMEM without phenol, RPMI with phenol, PBS, PBS with 10% FCS).

**Supplementary 5: Fully stained samples were unmixed using PBS+BSA controls with and without AF subtraction**. Fully stained samples in (**A** and **C**) PBS+BSA or (**B** and **D**) Pre-sort buffer were unmixed using PBS+BSA controls with (A and B) and without (**C** and **D**) autofluorescence subtraction.

**Supplementary Table 1:** List of fluorophores used in each experiment.

**Supplementary Table 2:** Instrument configurations: Detectors, lasers and bandpass filters for BD FACSDiscover™ S8, Thermo Fisher Invitrogen Bigfoot, Cytek Biosciences Aurora analyser and Cytek Biosciences Aurora CS system.

