## Supplementary figures and images for "The consequences of mismatched buffers in spectral cell sorting"

### Supplementary Figure 1

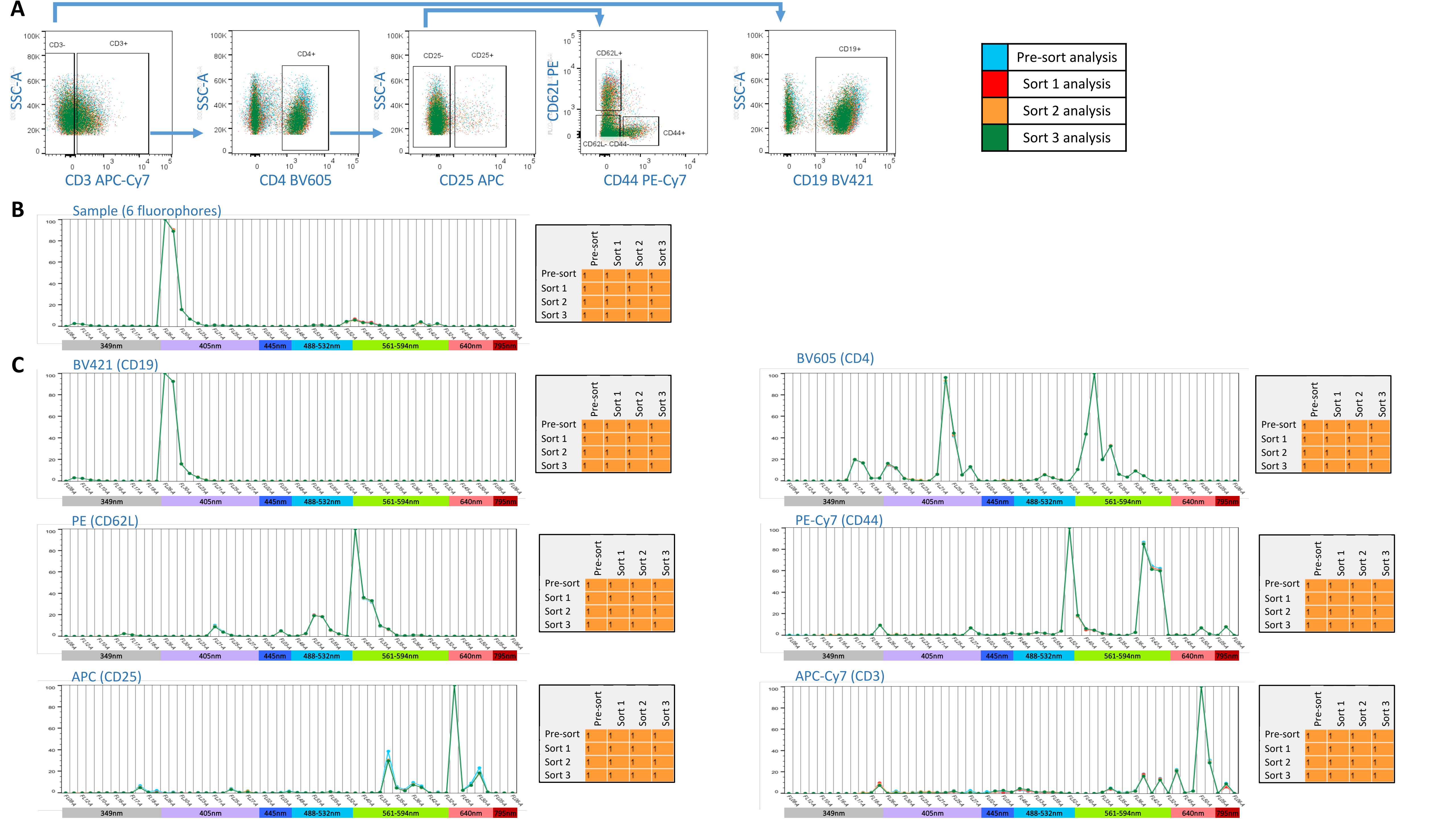

### Supplementary Figure 2

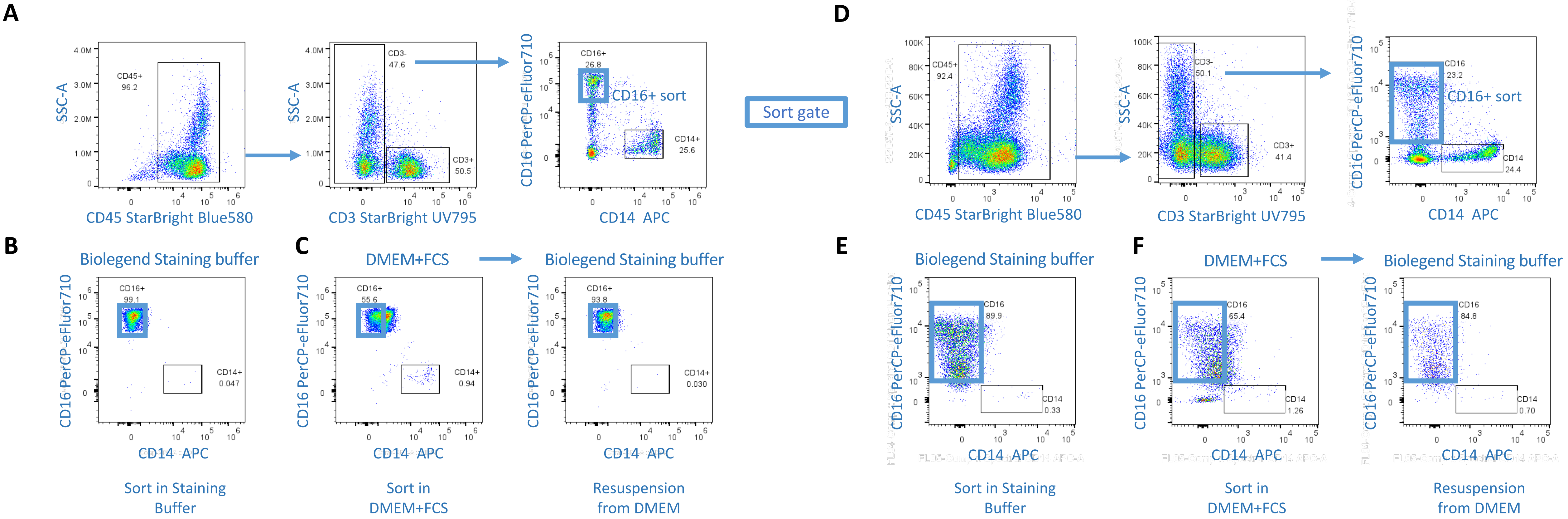

### Supplementary Figure 3

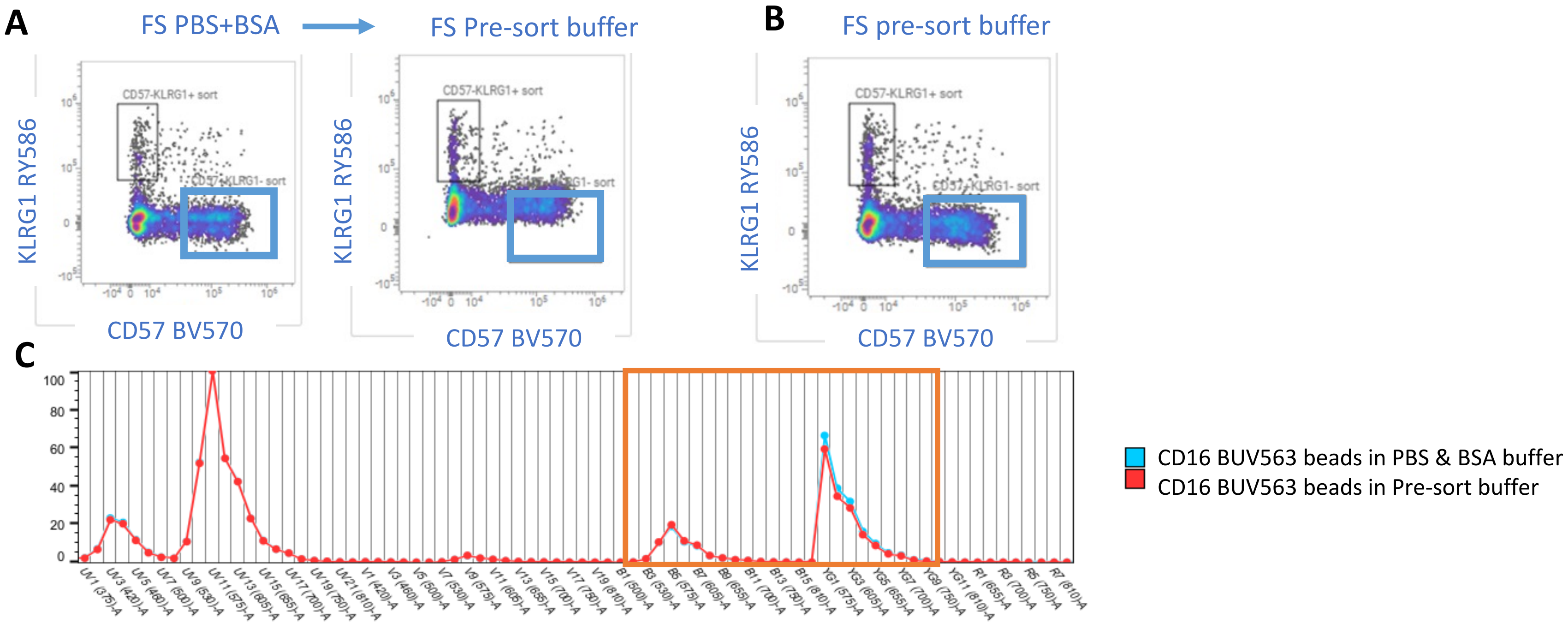

### Supplementary Figure 5

#### Autofluorescence figure

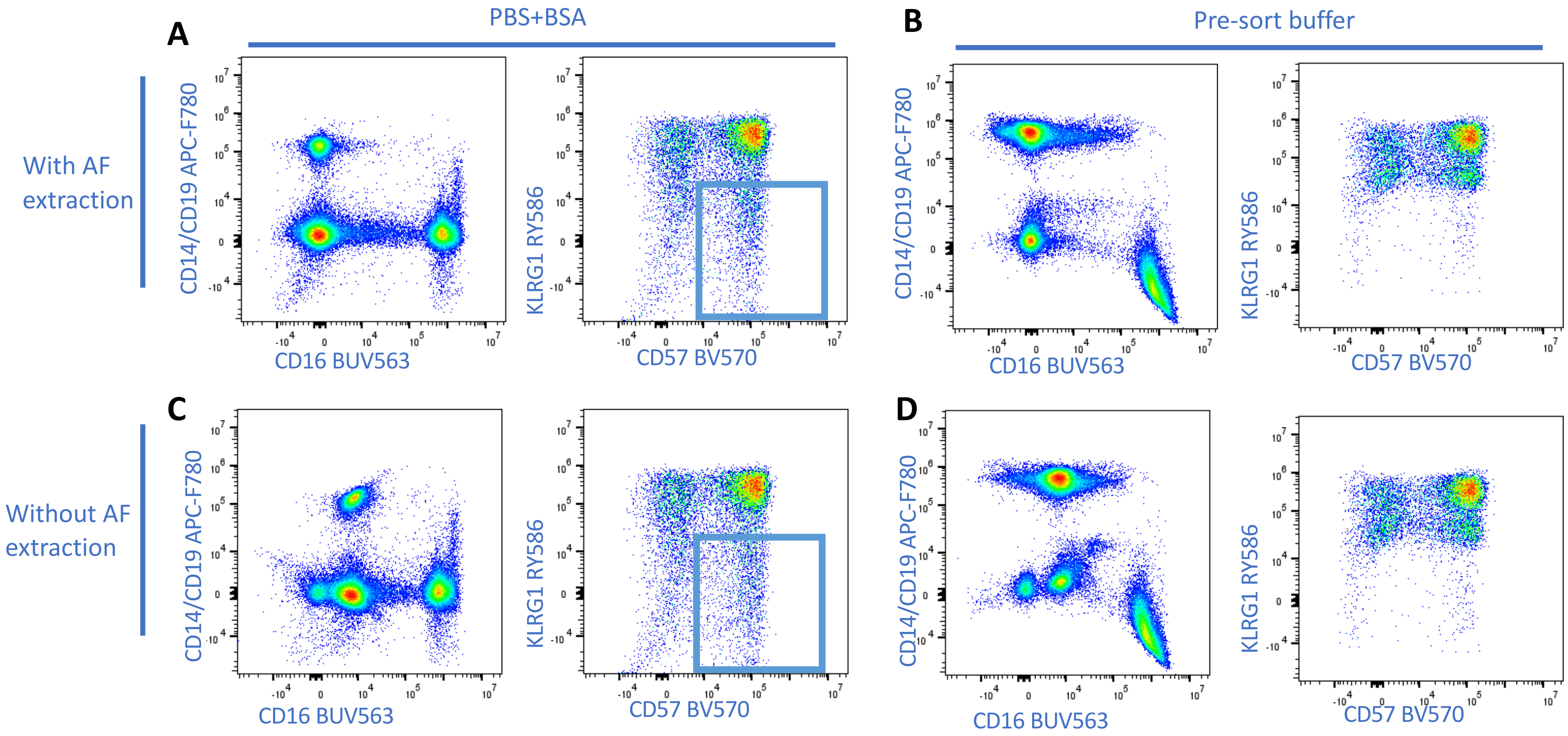
