## Supplementary Figure 4 for "The consequences of mismatched buffers in spectral cell sorting"

**A**

Lyophilized PBMCs (Veri cells ) - Lymphocytes

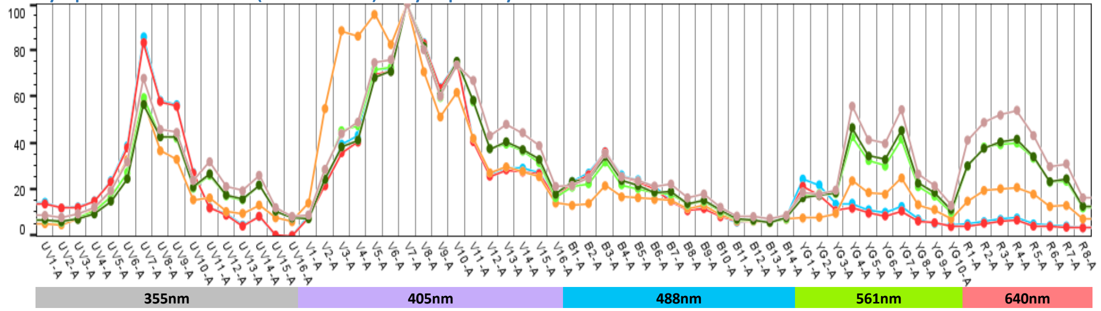


| | PBS | FACS Flow | DMEM w/o phenol | DMEM with phenol | RPMI with phenol | HANKS |
|---|---|---|---|---|---|---|
| PBS | 1 | 1 | 0.92 | 0.93 | 0.92 | 0.89 |
| FACS Flow | 1 | 1 | 0.92 | 0.92 | 0.91 | 0.89 |
| DMEM w/o | 0.92 | 0.92 | 1 | 0.94 | 0.92 | 0.91 |
| DMEM with | 0.93 | 0.92 | 0.94 | 1 | 1 | 0.99 |
| RPMI with | 0.92 | 0.91 | 0.92 | 1 | 1 | 1 |
| HANKS | 0.89 | 0.89 | 0.91 | 0.99 | 1 | 1 |

| | |
|---|---|
| (light blue) | PBS |
| (red) | BD FACS Flow |
| (orange) | DMEM w/o phenol and 10% FCS |
| (light green) | DMEM w/ phenol and 10% FCS |
| (dark green) | RPMI w/ phenol and 10% FCS |
| (pink-brown) | HANKS and 10% FCS |

**B**

Lyophilized PBMCs (Veri cells ) - Monocytes

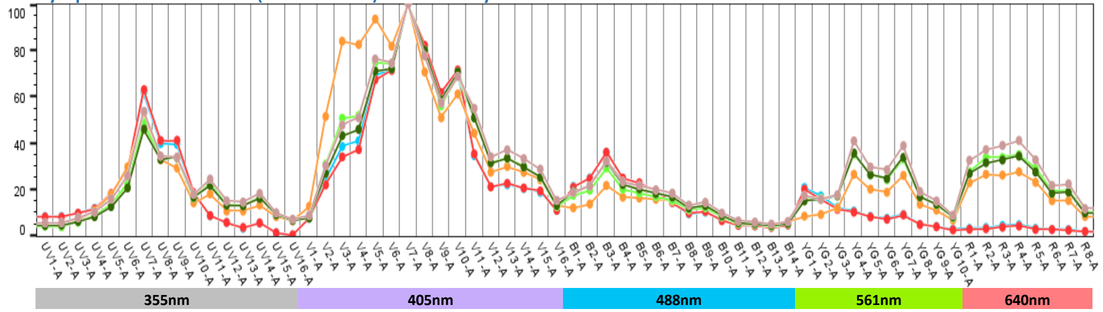


| | PBS | FACS Flow | DMEM w/o phenol | DMEM | RPMI | HANKS |
|---|---|---|---|---|---|---|
| PBS | 1 | 1 | 0.93 | 0.94 | 0.94 | 0.92 |
| FACS Flow | 1 | 1 | 0.92 | 0.93 | 0.93 | 0.92 |
| DMEM w/o | 0.93 | 0.92 | 1 | 0.97 | 0.96 | 0.96 |
| DMEM | 0.94 | 0.93 | 0.97 | 1 | 1 | 1 |
| RPMI | 0.94 | 0.93 | 0.96 | 1 | 1 | 1 |
| HANKS | 0.92 | 0.92 | 0.96 | 1 | 1 | 1 |

**C**

Frozen fixed Human PBMCs

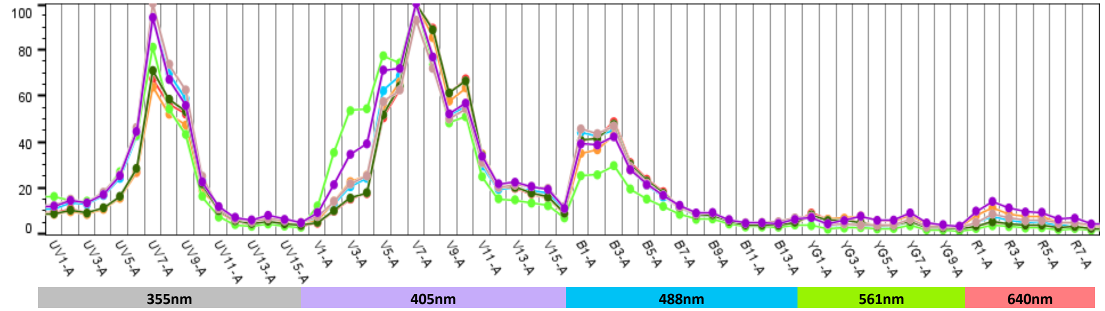


| | FACS Flow | DMEM | DMEM w FCS | DMEM w/o | RPMI | PBS | PBS w FCS |
|---|---|---|---|---|---|---|---|
| FACS Flow | 1 | 0.983727 | 0.983218 | 0.961221 | 0.987128 | 0.997854 | 0.993505 |
| DMEM | 0.983727 | 1 | 0.995383 | 0.937942 | 0.999717 | 0.975947 | 0.974027 |
| DMEM w | 0.983218 | 0.995383 | 1 | 0.955447 | 0.995651 | 0.974426 | 0.983607 |
| DMEM w/o | 0.961221 | 0.937942 | 0.955447 | 1 | 0.941861 | 0.953584 | 0.978269 |
| RPMI | 0.987128 | 0.999717 | 0.995651 | 0.941861 | 1 | 0.979916 | 0.977752 |
| PBS | 0.997854 | 0.975947 | 0.974426 | 0.953584 | 0.979916 | 1 | 0.991532 |
| PBS w FCS | 0.993505 | 0.974027 | 0.983607 | 0.978269 | 0.977752 | 0.991532 | 1 |

| | |
|---|---|
| (light blue) | BD FACS Flow |
| (red) | DMEM w/ phenol |
| (orange) | DMEM w/ phenol and 10% FCS |
| (light green) | DMEM w/o phenol |
| (dark green) | RPMI w/ phenol |
| (pink-brown) | PBS |
| (purple) | PBS with 10% FCS |

**D**

Frozen Mouse splenocytes

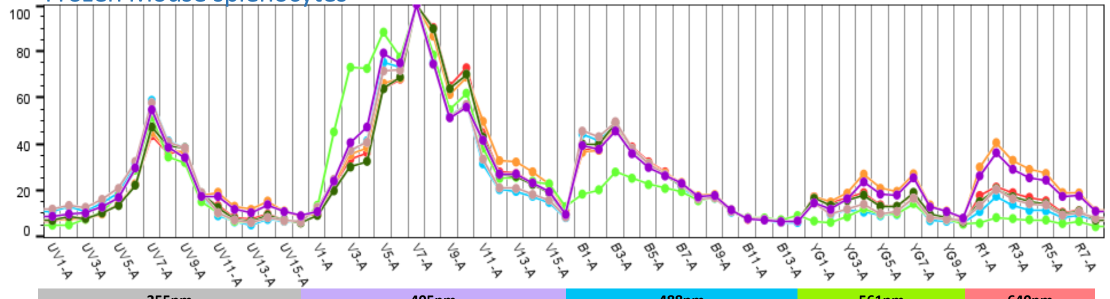


| | FACS Flow | DMEM | DMEM w FCS | DMEM w/o | RPMI | PBS | PBS w FCS |
|---|---|---|---|---|---|---|---|
| FACS Flow | 1 | 0.984081 | 0.965476 | 0.956222 | 0.987056 | 0.999099 | 0.981728 |
| DMEM | 0.984081 | 1 | 0.988409 | 0.945217 | 0.999125 | 0.986048 | 0.982785 |
| DMEM w | 0.965476 | 0.988409 | 1 | 0.926312 | 0.985754 | 0.971935 | 0.991411 |
| DMEM w/o | 0.956222 | 0.945217 | 0.926312 | 1 | 0.941348 | 0.953012 | 0.947867 |
| RPMI | 0.987056 | 0.999125 | 0.985754 | 0.941348 | 1 | 0.988467 | 0.981766 |
| PBS | 0.999099 | 0.986048 | 0.971935 | 0.953012 | 0.988467 | 1 | 0.986224 |
| PBS w FCS | 0.981728 | 0.982785 | 0.991411 | 0.947867 | 0.981766 | 0.986224 | 1 |
