## Supplementary Table 1 for "The consequences of mismatched buffers in spectral cell sorting"

| Figure | Experiment | Antibody for 1-10x10^6 cells | Fluorophore | Marker | Clone | Lot | Catalogue number | Manufacturer | Additional Buffers |
|---|---|---|---|---|---|---|---|---|---|
| Figures 1A, 2, Supplementary Figures 3 and 5 | Human 9c panel | 1:1000 dilution ratio | FVS440 | L/D | | 2207126 | 566332 | BD Biosciences | 10 µL Brilliant Staining Buffer Plus (BD Horizon #568264) |
| | | 2.5µL | BUV563 | CD16 | 3G8 | 3173922 | 748851 | BD Biosciences | |
| | | 2µL | V450 | CD7 | M-T701 | 3124066 | 642921 | BD Biosciences | |
| | | 1µL | BV480 | CD45RA | HI100 | 3135477 | 566114 | BD Biosciences | |
| | | 1µL | BV570 | CD57 | NK-1 | NA | NA (custom made) | BD Biosciences | |
| | | 5µL | RB545 | CD3 | UCHT1 | 3017835 | 569197 | BD Biosciences | |
| | | 5µL | RY586 | KLRG1 | Z7-205.rMAb | 2263584 | 568493 | BD Biosciences | |
| | | 2µL | Spark NIR685 | CD56 | 5.1H11 | B367292 | 362564 | Biolegend | |
| | | 5µL | APC-Fire810 | CD14 | 63D3 | B389915 | 367156 | Biolegend | |
| | | 1.25µL | APC-Fire810 | CD19 | HIB19 | B389981 | 302272 | Biolegend | |
| Figure 1B | Human 26c panel | 1.25µL | Brilliant Violet 421 | CD197 (CCR7) | G043H7 | B393745 | 353208 | Biolegend | 10 µL Brilliant Staining Buffer (BD Horizon #659611) 25 µL Human TruStain FcX™ (Biolegend #422301) 1X Tandem Stabilizer (Biolegend #421802) |
| | | 1.25µL | Pacific Blue | CD20 | 2H7 | B392751 | 302319 | Biolegend | |
| | | 1.25µL | Brilliant Violet 510 | IgM | MHM-88 | B354642 | 314522 | Biolegend | |
| | | 1.25µL | Brilliant Violet 570 | CD3 | UCHT1 | B411287 | 300436 | Biolegend | |
| | | 1.25µL | Brilliant Violet 605 | CD16 | 3G8 | B375535 | 302039 | Biolegend | |
| | | 1.25µL | Brilliant Violet 650 | CD28 | CD28.2 | B361779 | 302946 | Biolegend | |
| | | 1.25µL | Brilliant Violet 711 | CD38 | HIT2 | B399734 | 303528 | Biolegend | |
| | | 1.25µL | Brilliant Violet 750 | CD56 (NCAM) | 5.1H11 | B401476 | 362556 | Biolegend | |
| | | 1.25µL | Brilliant Violet 785 | CD279 (PD-1) | EH12.2H7 | B395800 | 329930 | Biolegend | |
| | | 1.25µL | Alexa Fluor® 488 | CD45RA | HI100 | B404214 | 304114 | Biolegend | |
| | | 1.25µL | Spark Blue™ 550 | CD14 | 63D3 | B314183 | 367147 | Biolegend | |
| | | 1.25µL | Spark Blue™ 574 | CD4 | SK3 | B413889 | 344679 | Biolegend | |
| | | 1.25µL | PerCP/Cyanine5.5 | CD8 | SK1 | B356155 | 344709 | Biolegend | |
| | | 1.25µL | PE | TCR γ/δ | B1 | B388480 | 331209 | Biolegend | |
| | | 1.25µL | Spark YG™ 581 | IgD | W18340F | B328367 | 307805 | Biolegend | |
| | | 1.25µL | PE/Dazzle™ 594 | CD141 (Thrombomodulin) | M80 | B354342 | 344119 | Biolegend | |
| | | 1.25µL | PE/Fire™ 700 | CD25 | M-A251 | B357964 | 356145 | Biolegend | |
| | | 1.25µL | PE/Cyanine7 | CD11c | 3.9 | B376429 | 301607 | Biolegend | |
| | | 1.25µL | PE/Fire™ 810 | HLA-DR | L243 | B382932 | 307683 | Biolegend | |
| | | 1.25µL | APC | CD127 (IL-7Rα) | A019D5 | B366605 | 351315 | Biolegend | |
| | | 1.25µL | Alexa Fluor® 647 | CD1c | L161 | B330200 | 331510 | Biolegend | |
| | | 1.25µL | Spark NIR™ 685 | CD19 | HIB19 | B411690 | 302269 | Biolegend | |
| | | 1.25µL | Alexa Fluor® 700 | CD123 | 6H6 | B374915 | 306039 | Biolegend | |
| | | 1:2000 dilution ratio | Zombie NIR™ | Fixable Viability Kit | Viability | B401652 | 423105 | Biolegend | |
| | | 1.25µL | APC/Fire™ 750 | CD45 | 2D1 | B374333 | 368517 | Biolegend | |
| | | 1.25µL | APC/Fire™ 810 | CD27 | O323 | B388988 | 302863 | Biolegend | |
| Figure 1C and Supplementary Figure 2 | Human 6c panel | 5µL | StarBright Blue 580 | CD45 | F10-89-4 | 100006954 | MCA87SBB580 | BioRad | 10 µL Super Bright Complete Staining Buffer (eBioscience #SB-4401-42) |
| | | 5µL | StarBright UV 795 | CD3 | UCHT1 | 10006205 | MCA463SBUV795 | BioRad | |
| | | 5µL | BV570 | CD4 | RPA-T4 | B341173 | 300533 | Biolegend | |
| | | 5µL | APC | CD14 | 61D3 | 2243488 | 17-0149-42 | Invitrogen | |
| | | 5µL | PerCP-eFluor710 | CD16 | 3G8 | 2338632 | 46-0149-42 | Invitrogen | |
| | | 5µL | SuperBright 702 | CD8 | RPA-TB | 2281536 | 67-0088-42 | Invitrogen | |
| Supplementary Figure 1 | Mouse 6c panel | 5µL | BV421 | CD19 | 6D5 | B212175 | 115549 | Biolegend | 10 µL Brilliant Staining Buffer (BD Horizon #659611) |
| | | 5µL | BV605 | CD4 | RM4-5 | B209226 | 100548 | Biolegend | |
| | | 5µL | PE | CD62L | MEL-14 | C0621060322502 | 50-0621 | Tonbo | |
| | | 5µL | PE-Cy7 | CD44 | IM7 | C0441060322602 | 60-0441 | Tonbo | |
| | | 5µL | APC | CD25 | PC61.5 | 2154046 | 102053 | Invitrogen | |
| | | 5µL | APC-Cy7 | CD3 | 17A2 | C0032060322252 | 25-0032 | Tonbo | |
