## Supplementary Table 2 for "The consequences of mismatched buffers in spectral cell sorting"

### BD FACSDiscover™ S8

| Detector name | Bandpass filter (laser wavelength (nm)) | Detector name | Bandpass filter (laser wavelength (nm)) | Detector name | Bandpass filter (laser wavelength (nm)) | Detector name | Bandpass filter (laser wavelength (nm)) | Detector name | Bandpass filter (laser wavelength (nm)) |
|---|---|---|---|---|---|---|---|---|---|
| UV1 (375) | 375/20 (349) | V1 (420) | 420/20 (405) | B1 (500) | 501/16 (488) | YG1 (575) | 575/14 (561) | R1 (655) | 655/20 (637) |
| UV2 (390) | 391/12.5 (349) | V2 (440) | 440/20 (420) | B2 (515) | 517/16 (488) | YG2 (590) | 590/16 (561) | R2 (675) | 675/20 (637) |
| UV3 (420) | 420/20 (349) | V3 (460) | 459/18 (420) | B3 (530) | 532/16 (488) | YG3 (605) | 607/18 (561) | R3 (700) | 698/26 (637) |
| UV4 (440) | 440/20 (349) | V4 (475) | 476/16 (420) | B4 (545) | 547/16 (488) | YG4 (625) | 623/14 (561) | R4 (725) | 724/26 (637) |
| UV5 (460) | 459/18 (349) | V5 (500) | 501/16 (420) | B5 (575) | 575/14 (488) | YG5 (655) | 655/20 (561) | R5 (750) | 752/30 (637) |
| UV6 (475) | 476/16 (349) | V6(515) | 517/16 (420) | B6 (590) | 590/16 (488) | YG6 (675) | 675/20 (561) | R6 (780) | 782/30 (637) |
| UV7 (500) | 501/16 (349) | V7 (530) | 532/16 (420) | B7 (605) | 607/18 (488) | YG7 (700) | 698/26 (561) | R7 (810) | 812/30 (637) |
| UV8 (515) | 517/16 (349) | V8 (545) | 547/16 (420) | B8 (625) | 623/14 (488) | YG8 (725) | 724/26 (561) | R8 (845) | 844/34 (637) |
| UV9 (530) | 532/16 (349) | V9(575) | 575/14 (420) | B9 (655) | 655/20 (488) | YG9 (750) | 752/30 (561) | | |
| UV10 (545) | 547/16 (349) | V10 (590) | 590/16 (420) | B10 (675) | 675/20 (488) | YG10 (780) | 782/30 (561) | | |
| UV11 (575) | 575/14 (349) | V11(605) | 607/18 (420) | B11 (700) | 698/26 (488) | YG11 (810) | 812/30 (561) | | |
| UV12 (590) | 590/16 (349) | V12(625) | 623/14 (420) | B12 (725) | 724/26 (488) | YG12 (845) | 844/34 (561) | | |
| UV13 (605) | 607/18 (349) | V13 (655) | 655/20 (420) | B13 (750) | 752/30 (488) | | | | |
| UV14 (625) | 623/14 (349) | V14(675) | 675/20 (420) | B14 (780) | 782/30 (488) | | | | |
| UV15 (655) | 655/20 (349) | V15 (700) | 698/26 (420) | B15 (810) | 812/30 (488) | | | | |
| UV16 (675) | 675/20 (349) | V16 (725) | 724/26 (420) | B16 (845) | 844/34 (488) | | | | |
| UV17 (700) | 698/26 (349) | V17 (750) | 752/30 (420) | | | | | | |
| UV18 (725) | 724/26 (349) | V18 (780) | 782/30 (420) | | | | | | |
| UV19 (750) | 752/30 (349) | V19 (810) | 812/30 (420) | | | | | | |
| UV20 (780) | 782/30 (349) | V20 (845) | 844/34 (420) | | | | | | |
| UV21 (810) | 812/30 (349) | | | | | | | | |
| UV22 (845) | 844/34 (349) | | | | | | | | |

### Thermo Fisher Invitrogen Bigfoot (6 & 9 laser)

| 6 and 9 laser | | 6 and 9 laser | | 9 laser | | 6 laser 488nm only. 9 laser 488/532 co-linear | | 6 laser 561nm only. 9 laser 561/532nm co-linear | | 6 and 9 laser | | 6 and 9 laser | |
|---|---|---|---|---|---|---|---|---|---|---|---|---|---|
| *Detector name* | *Bandpass filter (laser wavelength (nm))* | *Detector name* | *Bandpass filter (laser wavelength (nm))* | *Detector name* | *Bandpass filter (laser wavelength (nm))* | *Detector name* | *Bandpass filter (laser wavelength (nm))* | *Detector name* | *Bandpass filter (laser wavelength (nm))* | *Detector name* | *Bandpass filter (laser wavelength (nm))* | *Detector name* | *Bandpass filter (laser wavelength (nm))* |
| FL08-A | 387/11 (349) | FL26-A | 420/10 (405) | FL02-A | 465/22 (445) | FL48-A | 507/19 (488/532) | FL37-A | 575/15 (561/594) | FL49-A | 670/30 (640) | FL05-A | 800/12 (785) |
| FL13-A | 420/10 (349) | FL29-A | 434/17 (405) | FL00-A | 525/35 (445) | FL44-A | 549/15 (488/532) | FL40-A | 589/15 (561/594) | FL45-A | 700/13 (640) | FL04-A | 832/37 (785) |
| FL12-A | 434/17 (349) | FL30-A | 455/14 (405) | FL03-A | 583/30 (445) | FL53-A | 583/30 (488/532) | FL38-A | 605/15 (561/594) | FL51-A | 720/24 (640) | FL06-A | 860/LP (785) |
| FL14-A | 455/14 (349) | FL28-A | 473/15 (405) | FL01-A | 650LP (445) | FL47-A | 615/24 (488/532) | FL33-A | 625/15 (561/594) | FL50-A | 760/50 (640) | | |
| FL10-A | 473/15 (349) | FL23-A | 507/19 (405) | | | FL55-A | 670/30 (488/532) | FL39-A | 661/20 (561/594) | FL46-A | 770/LP (640) | | |
| FL11-A | 507/19 (349) | FL20-A | 549/15 (405) | | | FL54-A | 720/60 (488/532) | FL35-A | 685/15 (561/594) | | | | |
| FL16-A | 549/15 (349) | FL21-A | 575/15 (405) | | | FL52-A | 750/LP (488/532) | FL43-A | 700/13 (561/594) | | | | |
| FL19-A | 575/15 (349) | FL31-A | 615/24 (405) | | | | | FL36-A | 720/24 (561/594) | | | | |
| FL17-A | 615/24 (349) | FL25-A | 661/20 (405) | | | | | FL34-A | 760/50 (561/594) | | | | |
| FL15-A | 670/30 (349) | FL22-A | 710/20 (405) | | | | | FL42-A | 800/12 (561/594) | | | | |
| FL18-A | 728/40 (349) | FL27-A | 747/33 (405) | | | | | FL41-A | 832/37 (561/594) | | | | |
| FL09-A | 750/LP (349) | FL24-A | 770/LP (405) | | | | | FL32-A | 860/LP (561/594) | | | | |

### Cytek Biosciences Aurora Analyser / Aurora CS

| Detector name | Bandpass filter (laser wavelength (nm)) | Detector name | Bandpass filter (laser wavelength (nm)) | Detector name | Bandpass filter (laser wavelength (nm)) | Detector name | Bandpass filter (laser wavelength (nm)) | Detector name | Bandpass filter (laser wavelength (nm)) |
|---|---|---|---|---|---|---|---|---|---|
| UV1 | 372/15 (355) | V1 | 428/15 (405) | B1 | 508/20 (488) | YG1 | 577/20 (561) | R1 | 660/17 (640) |
| UV2 | 387/15 (355) | V2 | 443/15 (405) | B2 | 525/17 (488) | YG2 | 598/20 (561) | R2 | 678/18 (640) |
| UV3 | 427/15 (355) | V3 | 458/15 (405) | B3 | 542/17 (488) | YG3 | 615/20 (561) | R3 | 697/19 (640) |
| UV4 | 443/15 (355) | V4 | 473/15 (405) | B4 | 581/19 (488) | YG4 | 660/17 (561) | R4 | 717/20 (640) |
| UV5 | 458/15 (355) | V5 | 508/20 (405) | B5 | 598/20 (488) | YG5 | 678/18 (561) | R5 | 738/21 (640) |
| UV6 | 473/15 (355) | V6 | 525/17 (405) | B6 | 615/20 (488) | YG6 | 697/19 (561) | R6 | 760/23 (640) |
| UV7 | 514/28 (355) | V7 | 542/17 (405) | B7 | 660/17 (488) | YG7 | 720/29 (561) | R7 | 783/23 (640) |
| UV8 | 542/28 (355) | V8 | 581/19 (405) | B8 | 678/18 (488) | YG8 | 750/30 (561) | R8 | 812/34 (640) |
| UV9 | 581/31 (355) | V9 | 598/20 (405) | B9 | 697/19 (488) | YG9 | 780/30 (561) | | |
| UV10 | 612/31 (355) | V10 | 615/20 (405) | B10 | 717/20 (488) | YG10 | 812/34 (561) | | |
| UV11 | 664/27 (355) | V11 | 664/27 (405) | B11 | 738/21 (488) | | | | |
| UV12 | 691/28 (355) | V12 | 692/28 (405) | B12 | 760/23 (488) | | | | |
| UV13 | 720/29 (355) | V13 | 720/29 (405) | B13 | 783/23 (488) | | | | |
| UV14 | 750/30 (355) | V14 | 750/30 (405) | B14 | 812/34 (488) | | | | |
| UV15 | 780/30 (355) | V15 | 780/30 (405) | | | | | | |
| UV16 | 812/34 (355) | V16 | 812/34 (405) | | | | | | |
